# Deletion of cytoplasmic β/γ-actin in the mouse heart protects from disease by augmenting sarcolemma stability

**DOI:** 10.64898/2026.07.24.740593

**Authors:** Yasuhide Kuwabara, Caitlin Keezer, Eaman Abay, Suh-Chin J. Lin, Jeffery D. Molkentin

## Abstract

The actin cytoskeleton of the cardiomyocyte is organized into two structurally and functionally distinct filament networks. The sarcomeric thin filaments, built primarily from cardiac α-actin (αCA), generate contractile force, whereas a separate subsarcolemmal cytoplasmic network, built from β-actin (*Actb* gene) and γ-actin (*Actg1* gene), lies beneath the sarcolemma and is comparatively understudied in the mature heart. We hypothesized that this cytoplasmic actin network is required for sarcolemmal membrane integrity, signal transduction, and mechanosensing in the adult cardiomyocyte, and we generated cardiomyocyte-specific *Actb* and *Actg1* double-gene deleted mice, using loxP (fl)-targeted *Actb*^fl/fl^ and *Actg1*^fl/fl^ alleles combined with an αMHC (Myh6) promoter-driven Cre-recombinase transgene, to test this directly. Deletion of cytoplasmic β-actin and γ-actin from cardiomyocytes (*Actb/g1^fl/fl-Myh6-Cre^* mice) drove compensatory upregulation of a γ-interferon like stress response with compensatory upregulation of skeletal α-actin (αSKA) and smooth muscle α-actin (αSMA) protein in adult cardiomyocytes, without altering baseline cardiac structure or function. To test if lost β-actin and γ-actin in the mouse heart impacts sarcolemmal stability, we crossed *Actb/g1^fl/fl-Myh6-Cre^*mice onto the dystrophin-deficient *mdx* background, which is characterized by a fragile sarcolemma. Unexpectedly, hearts from *Actb/g1^fl/fl-Myh6-Cre^*; *mdx* mice showed greater membrane stability than *mdx* hearts alone with intact *Actb*/*Actg1*. We also observed that *Actb/g1^fl/fl-Myh6-Cre^*mice subjected to chronic pressure overload by transverse aortic constriction (TAC) were protected and developed less cardiac hypertrophy, had better preserved systolic function, and improved survival. Together, these data indicate induction of the cytoplasmic β-actin/γ-actin network in the heart during disease stimulation is maladaptive and weakens the sarcolemma, and 2 downstream mechanisms are considered that could mediate this effect.

## Introduction

Heart failure remains one of the most prevalent and costly chronic diseases in the United States, affecting more than 6 million American adults and contributing to roughly one in eight deaths [1]. Once initiated, heart failure typically follows a progressive and largely self-reinforcing course of ventricular remodeling, marked by cardiomyocyte hypertrophy, interstitial fibrosis, and a gradual decline in contractile performance. A defining molecular feature of this remodeling process is the reactivation of a “fetal gene program,” in which the stressed adult cardiomyocyte re-expresses structural, contractile, and stress-responsive genes that are otherwise restricted to fetal development and silenced soon after birth [2]. This program includes several sarcomeric and cytoskeletal isoform switches, and while some elements of fetal gene re-expression appear to represent compensatory attempts by the myocyte to withstand injury, others are likely to contribute to disease progression itself.

A key structural determinant of cardiomyocyte survival under mechanical or hemodynamic stress is the integrity of the sarcolemma, the plasma membrane that must simultaneously withstand the repetitive mechanical load of contraction and serve as the principal structural interface between the cardiomyocyte cytoskeleton and the extracellular matrix. Weakening of the sarcolemma is a well-established contributor to cardiac disease pathogenesis, most clearly illustrated in the *mdx* mouse model of Duchenne muscular dystrophy, in which loss of the cytoskeletal protein dystrophin destabilizes the costameric lattice that anchors the sarcolemma, rendering the membrane fragile and prone to injury during contraction [3,4]. With a fragile sarcolemma, mechanical stress produces microscopic membrane tears that permit uncontrolled extracellular calcium influx, which in turn activates calcium-dependent proteases and drives cardiomyocyte and skeletal muscle myofiber necrosis [5,6]. Conversely, cardiomyocyte-specific overexpression of α7β1 integrin, a principal costameric receptor that links the cortical cytoskeleton to the extracellular matrix, protects the heart from acute ischemia-reperfusion injury induced heart dysfunction [7]. Similarly, overexpression of thrombospondin-4 in the mouse heart strengthens the sarcolemma by augmenting integrins, which protected from heart failure following chronic pressure overload or ischemia/reperfusion injury [8]. Thrombospondin-4 overexpression in skeletal muscle similarly protected mouse models of muscular dystrophy from worsening disease [9]. Collectively these results demonstrate that reinforcing costameric and sarcolemmal architecture, whether through native adaptation or genetic manipulation, is cardioprotective.

The mammalian genome encodes six actin isoforms that, despite differing from one another by only 4 amino acids, are segregated into distinct subcellular compartments with largely non-redundant functions [10,11]. In the heart, actin filaments are organized into two principal networks. The first is the sarcomeric thin filament network, which during development is composed of three α-actin isoforms, cardiac (encoded by *Actc1*), skeletal (*Acta1*), and smooth muscle (*Acta2*). *Acta1* and *Acta2* are progressively silenced in the heart over postnatal development, such that the mature adult sarcomere is composed almost exclusively of cardiac α-actin (*Actc1* gene), although with disease stimulation compensatory re-expression of *Acta1* and *Acta2* is observed [12]. The second network is a subsarcolemmal cytoplasmic actin scaffold built from cytoplasmic β- and γ-actin, encoded by *Actb* and *Actg1*, which in non-muscle cells regulates the assembly, turnover, and recycling of focal adhesion and integrin complexes [10]. As with the sarcomeric network, *Actb* and *Actg1* expression declines substantially over heart development, so that the cytoplasmic actin network is not a prominent structure in the unstressed, mature cardiomyocyte. However, acute and chronic cardiac disease induces greater *Actb* and *Actg1* expression [13], together with more modest re-expression of *Acta1* and *Acta2* [14]. Despite these consistent observations across multiple disease models, whether re-expression of the cytoplasmic actin network itself, or the coincident re-expression of sarcomeric skeletal and smooth muscle α-actin isoforms is beneficial or harmful to the diseased heart has not been established.

Functional dissection of individual actin isoforms has been pursued in skeletal muscle using conditional, Cre-recombinase-mediated gene targeting. Muscle-specific deletion of *Actg1* produces a progressive myopathy marked by myofiber necrosis, regeneration, and disorganization of the costameres that couple myofibrils to the sarcolemma, despite otherwise normal muscle development [15]. Skeletal muscle-specific ablation of *Actb* similarly produces a progressive quadriceps myopathy [16]. Notably, when the conditional *Actg1* allele was combined with the dystrophin-deficient *mdx* background, loss of cytoplasmic γ-actin did not exacerbate the membrane instability and myofiber necrosis already present in *mdx* muscle, indicating that cytoplasmic actin and dystrophin may function, at least in part, within a shared pathway that maintains sarcolemmal integrity [17]. These skeletal muscle studies establish that the cytoplasmic actin isoforms are required for normal costameric and membrane architecture, but how *Actb* and *Actg1* function in the heart when re-expressed with disease stimulation is unknown.

Here we deleted *Actb* and *Actg1* from the mouse heart using a cardiomyocyte-specific Cre transgene approach to directly test the functional importance of the cytoplasmic actin network in the postnatal and adult heart. In contrast to the myopathic phenotype produced by cytoplasmic actin deletion in skeletal muscle, we found that cardiomyocyte-specific loss of β-actin and γ-actin was well tolerated at baseline but triggered compensatory upregulation of αSKA and αSMA at the cardiomyocyte sarcolemma. This compensatory response was associated with increased resistance to membrane instability when combined with the *mdx* mutation, and it improved survival and reduced cardiac hypertrophy following chronic pressure overload. These data suggest that re-expression of a more prominent cytoplasmic actin network is maladaptive in the heart, or that compensatory induction of skeletal muscle α-actin and smooth muscle α-actin isoforms is protective to the heart.

## Materials and Methods

### Animals

*Actb* and *Actg1* loxP site targeted mice are kindly gifted from Dr. James M. Ervasti at University of Minnesota [18,19]. In this article, two transgenic mouse lines with the α-myosin heavy chain (Myh6) promoter directing Cre recombinase (Myh6-Cre Tg) in the C57BL/6 background were utilized [20,21]. Both transgenic lines are also at the Jackson Laboratory (#009074: Tg(Myh6-cre)1Jmk/J and #011038: B6.FVB-Tg(Myh6-cre)2182Mds/J). Tg(Myh6-cre)1Jmk/J line was basically used except for the *mdx* crossing experiments (**Figure 3** and **Supplementary Figure 1**) because both the *mdx* mutant allele and the Tg(Myh6-cre)1Jmk/J Myh6-Cre Tg line is X-linked, which is also why only male mice were used. As control mice of *Actb/g1^fl/fl-Myh6-Cre^* mice, male *Actb/g1^+/+-Myh6-Cre^*mice were used. Inducible cardiomyocyte-specific *Actb* and *Actg1* bicistronic dual-transgenic mice were originally generated with tetracycline-off system in the C57BL/6J background [22]. Internal ribosome entry site (IRES) was introduced between *Actb* and *Actg1* for *Actg1* expression. To attempt proper subcellular distribution of β-actin, a 60 nucleotide long sequence termed the zip code was inserted after the *Actb* stop codon. The Myh6 promoter-driven tetracycline-controlled transactivator (tTA) Tg mice were described previously [22] and maintained in the laboratory in the C57BL/6 background. Mice were never given doxycycline or tetracycline, so the double transgenic mice were constitutively in the induced state of expression.

### Transverse Aortic Constriction Surgery

Transverse Aortic Constriction (TAC) surgery was conducted as previously reported [20,23]. Briefly, eight-week-old mice were anesthetized by 3% isoflurane and intubated with an 18-gauge catheter. Anesthesia was maintained during surgery with a mouse ventilator delivering 1.7% isoflurane. A thoracotomy was performed, and the transverse aorta was isolated and tied around a 27-gauge needle, which was then removed to generate the defined constriction. After extubation, sustained-release buprenorphine (0.2 mg/kg) was injected subcutaneously in a volume of 0.03 ml to reduce pain, and mice were monitored daily after surgery. Sham surgery was performed identically, except that the aorta was not constricted.

### Animal Welfare and Ethics

Animals were handled in accordance with the principles and procedures of the Guide for the Care and Use of Laboratory Animals. All proposed procedures were approved by the Institutional Animal Care and Use Committee (IACUC) at Cincinnati Children’s Hospital Medical Center (CCHMC) in the USA. The IACUC ID is 2024-0043 and the approval date is Sep/4/2024. Animal groups and experiments were handled in a blinded manner where possible. No mice were excluded during the experiments. Randomization was not performed given that all mice were of the same genotypes and identical strain, and only age-matched littermates were compared. ARRIVE guidelines were followed in all mouse experimentation [24]. No human materials or subjects were used. All mice were housed in a germ-free barrier environment with free access to food and water on a 14-hour day/10-hour night cycle and were observed daily by veterinary staff. Pain management in mice involved sustained-release buprenorphine (0.2 mg/kg) injected subcutaneously in a volume of 0.03 ml.

### Echocardiography

Echocardiography was performed as previously described [20]. Briefly, mice were anesthetized with 3% isoflurane for induction, which was maintained at 1.7% isoflurane during imaging. Images were recorded on the ventral side of the chest in a blinded manner using a Vevo 3100 (Fujifilm VisualSonics) and analyzed with Vevo Lab software (Fujifilm VisualSonics). Parameters including ejection fraction (EF) were determined from M-mode images, with analyses also conducted in a blinded manner.

### Survival Analysis

Animal care technicians, under the supervision of two veterinarians, monitored the mice daily and assessed well-being according to the IACUC-approved guidelines for this study in a blinded manner, as well as monitoring for survival and achieving an end point without waiting for mortality [25]. For example, mice deemed excessively moribund were first treated with gel diet placed at the bottom of the cage to provide hydration and nutrition; if mice progressed and were no longer able to acquire nutrients or hydration, they were removed from the study and euthanized (considered a death event in the survival studies), as approved within the IACUC protocol and by the Office of Laboratory Animal Welfare.

### Adult Cardiomyocyte Isolation

Adult cardiomyocytes were isolated using the retrograde Langendorff perfusion method as previously described, with some modifications [23]. Briefly, mice were euthanized and hearts rapidly excised, rinsed in 1× phosphate-buffered saline (PBS), and the aorta cannulated and perfused with perfusion buffer on a Langendorff apparatus. The perfusion buffer (pH 7.4) consisted of 113 mM NaCl, 4.7 mM KCl, 0.6 mM KH_2_PO_4_, 0.6 mM Na_2_HPO_4_, 1.2 mM MgSO_4_·7H_2_O, 12 mM NaHCO_3_, 10 mM KHCO_3_, 0.032 mM phenol red, 10 mM HEPES, 30 mM taurine, 10 mM 2,3-butanedione monoxime (BDM), and 5.5 mM glucose. Hearts were digested with buffer containing collagenase II (#LS004177, Worthington) and 40 μM CaCl_2_. After digestion, the atrium was removed and the ventricle triturated, filtered through 240 μm mesh, and combined with stopping buffer containing 10% fetal bovine serum and 12.5 μM CaCl_2_. Isolated cardiomyocytes were allowed to settle by gravity and washed twice with stopping buffer. After the final settling step, the cardiomyocyte pellet was snap-frozen in liquid nitrogen for Western blotting.

### Protein Extraction

Protein extraction from heart tissue or isolated cardiomyocytes in 8–10-week-old mice was conducted as previously described [20,25]. Briefly, cell pellets or tissue samples were homogenized in 10 mM Tris (pH 7.5), 150 mM NaCl, 4% glycerol, 0.5 mM sodium metabisulfite, 1% Triton X-100, 0.05% SDS, and 0.1% sodium deoxycholate, with complete mini proteinase inhibitor cocktail (#04693124001, Roche) added just before homogenization. Homogenates were centrifuged at 14,000 × g for 15 min at 4°C, and the supernatant was snap-frozen and stored at -80°C until use.

Membrane fraction isolation from heart tissue was carried out as previously described [9,26]. Briefly, heart tissue was homogenized in lysis buffer (20 mM Na_4_P_2_O_7_, 20 mM NaH_2_PO_4_, 1 mM MgCl_2_, 0.3 M sucrose, 0.5 mM EDTA, pH 7.1) with proteinase inhibitor cocktail added just before homogenization. Homogenates were centrifuged at 14,000 × g for 15 min at 4°C; the supernatant was retained for ultracentrifugation and the pellet re-homogenized. After a second centrifugation at 14,000 × g for 15 min at 4°C, the combined supernatant was ultracentrifuged at 30,000 × g for 30 min at 4°C, and the resulting pellet was snap-frozen and stored at -80°C until use.

The sarcomere fraction was isolated as previously described, with modifications [27]. Heart tissue was homogenized in F-60 buffer (pH 7.0; 60 mM KCl, 30 mM imidazole, 2 mM MgCl_2_) with proteinase inhibitor cocktail, centrifuged at 12,000 × g for 10 min at 4°C, and the supernatant discarded; this wash step was repeated three times. Urea buffer (4 M urea, 50 mM Tris-HCl, 1 M thiourea, 0.4% CHAPS, pH 7.5) with proteinase inhibitor cocktail was then added to the final pellet. After homogenization and sonication, samples were centrifuged at 15,300 × g for 10 min at 4°C, and supernatants were stored at - 80°C until use.

### Western Blotting

Protein concentrations were determined using the DC protein assay kit (#5000112, Bio-Rad) or Protein assay kit (#5000001, Bio-Rad). Samples were prepared with 5× SDS loading dye, boiled at 100°C for 10 min, and subjected to polyacrylamide gel electrophoresis (PAGE). Gels were transferred to Immobilon-P PVDF membranes (#IPVH00010, Millipore Sigma), blocked with 5% milk protein in 1× PBS for 1 hour, and incubated with primary antibody in 5% milk protein/Tris-buffered saline with 0.1% Tween 20 (TBST) at 4°C overnight. Membranes were washed with TBST and incubated with secondary antibody (IRDye, LI-COR, 1:3000) in 5% milk protein/TBST with 0.01% SDS at room temperature for 1 hour. After 3 washes with TBST, membranes were scanned on an Odyssey CLx imaging system with Image Studio software (LI-COR).

Primary antibodies used for Western blots were anti-β-actin (#A1978, Sigma-Aldrich, 1:1000), anti-γ-actin (#A8481, Sigma-Aldrich, 1:1000), anti-pan-actin (#NBP2-25142, Novus Biologicals, 1:1000), anti-phospho ERK1/2 (#9101, Cell Signaling, 1:1000), anti-ERK1/2 (#9102, Cell Signaling, 1:1000), anti-MEK (#9122, Cell Signaling, 1:1000), anti-cardiac α-actin (#NBP2-61474, Novus Biologicals, 1:1000), anti-skeletal α-actin (#17521-1-AP, Proteintech, 1:1000), anti-smooth muscle α-actin (αSMA) (#A2547, Sigma-Aldrich, 1:1000), γSMA (#ab123034, Abcam, 1:1000), anti-α-actinin-2 (#A7811, Sigma-Aldrich, 1:1000), anti-α-actinin-1 (#ab68194, Abcam, 1:1000), anti-α-actinin-4 (#ab108198, Abcam, 1:1000), anti-integrin β1 (#MAB1900, Merck Millipore, 1:500), and anti-GAPDH (#10R-G109A, Fitzgerald, 1:5000).

### RNA isolation, bulk RNA sequencing, and bioinformatic analysis

Total RNA was isolated with TRIzol reagent (#15596018, Thermo Fisher Scientific) from heart tissue as previously described [25,28]. RNA quality was assessed using the RNA 6000 Nano Assay (#5067-1511, Agilent). Library preparation and Illumina platform bulk RNA sequencing were carried out by Illumina Inc. using a NovaSeq 6000. Bioinformatic analysis was performed to identify differentially expressed genes (DEGs) in hearts of *Actb/g1^+/+-Myh6-Cre^* mice versus *Actb/g1^fl/fl-Myh6-Cre^* mice at 2 weeks of age. RNA-seq data sets are deposited in the Gene Expression Omnibus (GEO) repository under accession number GSE339442 (embargoed until publication acceptance). A volcano plot was generated with GraphPad Prism 10 (GraphPad Software) by plotting log2 fold-change (log2FC) against –log10(adjusted p-value), with genes classified as upregulated, downregulated, or not significant according to adjusted p-value < 0.05 and |log2FC| > 1; horizontal and vertical dashed lines indicate these cutoffs. Gene Ontology enrichment analysis was performed on the subset of 62 upregulated genes (adjusted p-value < 0.05, log2FC > 2) using the Database for Annotation, Visualization, and Integrated Discovery (DAVID).

### Immunohistochemistry

Immunohistochemistry was conducted as previously described [20]. Mouse hearts were perfused with relaxing buffer for 5 min, followed by perfusion fixation with 4% paraformaldehyde (PFA) for 5 min, then excised and immersed in 4% PFA for 5 hours at 4°C. After three washes in 1× PBS, tissue was embedded in O.C.T. compound (#4583, Tissue-Tek) and cut into 7 µm sections on a cryostat (#CM1860, Leica) at -20°C. Sections were washed three times in 1× PBS, permeabilized in 0.1% Triton X-100/1× PBS for 10 min, blocked with 5% goat serum in 1× PBS for 15 min, and incubated overnight with primary antibodies: anti-skeletal α-actin (#17521-1-AP, Proteintech, 1:100), anti-α-actinin-2 (#A7811, Sigma-Aldrich, 1:300), anti-smooth muscle α-actin (#A2547, Sigma-Aldrich, 1:200), anti-TMOD1 (#NBP2-00955, Novus Biologicals, 1:200), and anti-β-actin (#A1978, Sigma-Aldrich, 1:100). The next day, slides were washed three times with 1× PBS and incubated with secondary antibodies (1:400): Alexa Fluor 568 goat anti-rabbit IgG (#A-11011, Invitrogen), Alexa Fluor 488 goat anti-mouse IgG1 (#A-21121, Invitrogen), Alexa Fluor 647 goat anti-mouse IgG1 (#A-21240, Invitrogen), and Alexa Fluor 647 goat anti-mouse IgG2b (#A-21242, Invitrogen). Invitrogen A-21131 (AF405) was used for anti-mouse IgG2a. For membrane staining, Alexa Fluor 488-conjugated wheat germ agglutinin (WGA) (#W11261, Thermo Fisher Scientific) was included with the secondary antibody; for sarcomeric actin thin filament staining, Alexa Fluor 488-conjugated phalloidin (#A12379, Invitrogen) was similarly included. After 1 hour of incubation at room temperature, slides were washed three times with 1× PBS and mounted using ProLong Diamond antifade mountant with DAPI (#P36962, Thermo Fisher Scientific). Images were captured on an A1 confocal microscope or AX/AX R with NSPARC microscope and analyzed with NIS-Elements software (Nikon).

### Evans blue dye uptake experiment in vivo

Evans blue dye (EBD) (#E2129, Sigma-Aldrich) at 100 mg/kg body weight was injected intraperitoneally in 100 μl PBS, and 24 hours later the heart was excised [25,26]. After washing in ice-cold PBS, hearts were embedded in O.C.T. compound (#4583, Tissue-Tek) and cut into 7 µm sections on a cryostat (#CM1860, Leica). For membrane staining, Alexa Fluor 488-conjugated WGA (#W11261, Thermo Fisher Scientific) staining was performed for 1 hour at room temperature. Images were captured on an A1 confocal microscope with NIS-Elements software (Nikon), and EBD-positive cardiomyocytes were identified directly by the Texas Red channel.

### Statistics

For comparisons of two groups, we first assessed whether data were normally distributed unless otherwise noted. Normally distributed data were analyzed by unpaired two-tailed t-tests; non-normally distributed data were analyzed by the Mann-Whitney test. For more than two groups, one-way analysis of variance (ANOVA) with Tukey’s post hoc test was performed. Survival was analyzed by log-rank test. A P value less than 0.05 was considered statistically significant. All statistical analyses were performed using GraphPad Prism 10 (GraphPad Software) unless otherwise noted.

## Results

### Cardiomyocyte-specific deletion of the cytoplasmic actin network is well tolerated at baseline

To determine the functional role of the cytoplasmic actin network in the adult mouse heart, we used conditional gene-targeted mice for *Actb* and *Actg1*, which were previously loxP-targeted and shown to undergo robust, Cre-dependent gene deletion [18,19]. We crossed these *Actb* and *Actg1* loxP-targeted mice with the αMHC (Myh6) promoter-driven Cre recombinase transgenic line to generate cardiomyocyte-specific *Actb* and *Actg1* double-knockout mice (*Actb/g1^fl/fl-Myh6-Cre^*, or B/G1-CKO) (**Figure 1A**). This Myh6-Cre driver produces early gene deletion in the developing heart, with Cre activity continuing through postnatal and adult stages [29]. At 16 weeks of age, we compared β- and γ-actin protein levels in purified cardiomyocytes isolated from B/G1-CKO hearts and littermate controls (**Figure 1B**) and found a dramatic loss of expression of each isoform. Total cardiomyocyte actin content was unchanged, reflecting the overwhelming abundance of sarcomeric α-actin relative to the cytoplasmic pool, and phospho-ERK1/2 and MEK1/2 levels were similarly unaffected (**Figure 1C**). Despite this loss of the cytoplasmic actin network throughout development and into adulthood, B/G1-CKO mice showed no overt disease phenotype: cardiac function by echocardiography and heart weight-to-body weight ratio were indistinguishable from controls (**Figure 1D** and **1E**), indicating that viable B/G1-CKO mice are phenotypically normal at baseline.

**Figure 1.**
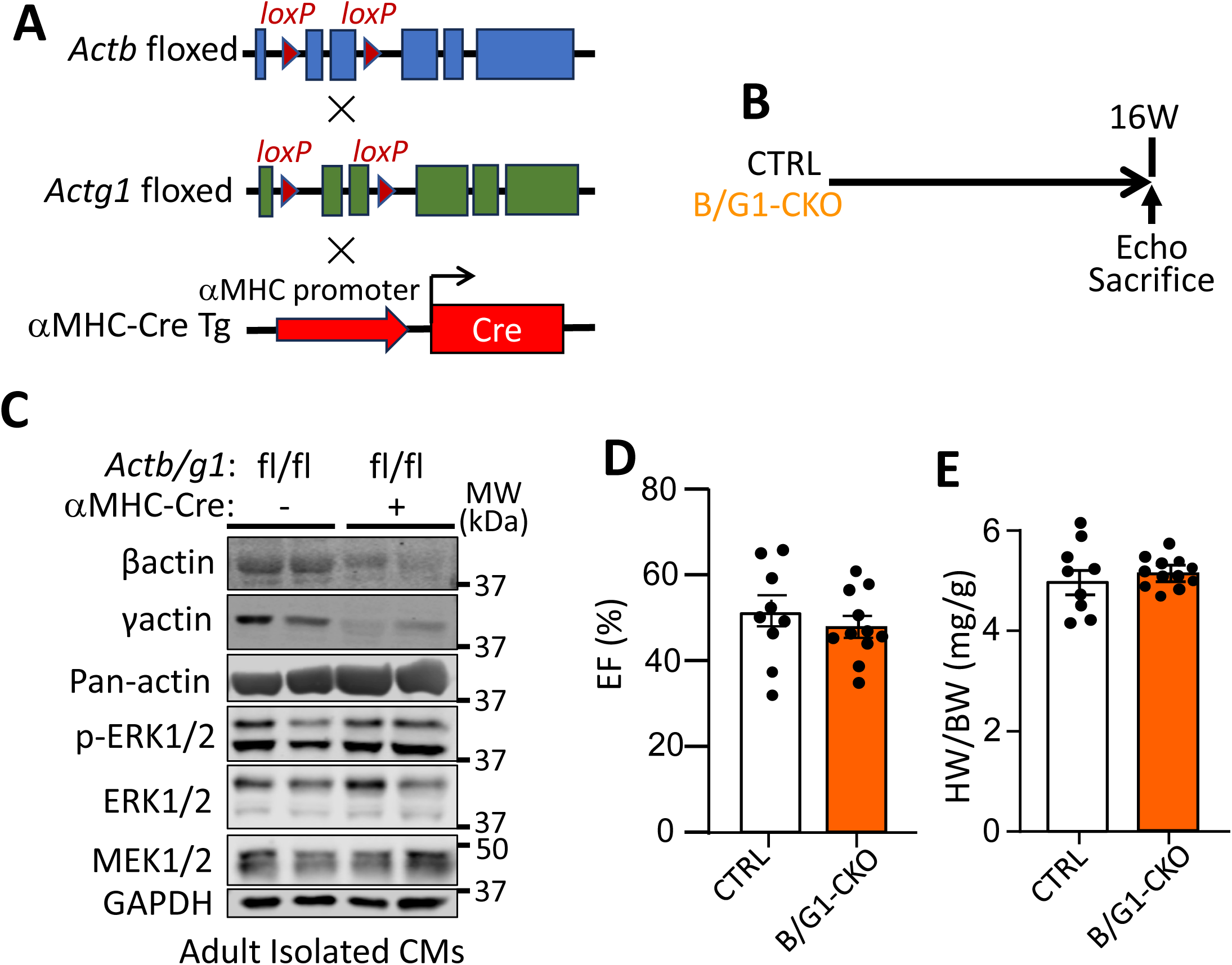
Cardiomyocyte-specific *Actb* and *Actg1* gene deletion in mice. **(A)** Cardiomyocyte-specific *Actb* and *Actg1* double-gene deleted mice (*Actb/g1^fl/fl-Myh6-Cre^*: B/G1-CKO) were generated by crossing loxP-targeted *Actb* and *Actg1* mice with αMHC (Myh6) promoter-driven Cre recombinase transgenic mice. **(B)** Experimental time-line to characterize the cardiac phenotype of B/G1-CKO mice; *Actb/g1^+/+-Myh6-Cre^* mice served as controls (CTRL) unless otherwise noted, and mice were analyzed at 16 weeks of age (16W) by echocardiography (Echo). **(C)** Western blots of β-actin, γ-actin, pan-actin, phospho-ERK1/2, total ERK1/2, and MEK1/2 in protein extracts from isolated adult cardiomyocytes of B/G1-CKO hearts. GAPDH served as a loading control. Numbers indicate molecular weight (kDa). **(D)** Ejection fraction percentage (EF%) by echocardiography at 16 weeks of age (n = 9 control, n = 11 B/G1-CKO). **(E)** Heart weight/body weight ratio (HW/BW) at 16 weeks of age (n = 9 control, n = 11 B/G1-CKO). Data in D and E are shown as scatter plots with bars indicating mean ± SEM.

### Loss of cytoplasmic actin triggers an innate immune transcriptional signature and compensatory α-actin induction

To further investigate potential pathologic consequences of deleting the cytoplasmic actin network, we performed bulk RNA sequencing on B/G1-CKO hearts versus controls and identified only 62 upregulated and 40 downregulated genes as significantly changed (**Figure 2A, Supplementary Tables 1 and 2**). Gene Ontology Biological Process (GO-BP) enrichment analysis of the upregulated genes identified categories related predominantly to the interferon response and innate immune activation, pathways also implicated in membrane regulation during viral entry and host-pathogen sensing (**Figure 2B**). This finding suggests that loss of the cytoplasmic actin network from the developing and adult heart is sensed by the cardiomyocyte in a manner resembling a change in sarcolemmal membrane dynamics, similar to signaling elicited by viral or pathogen challenge.

**Figure 2.**
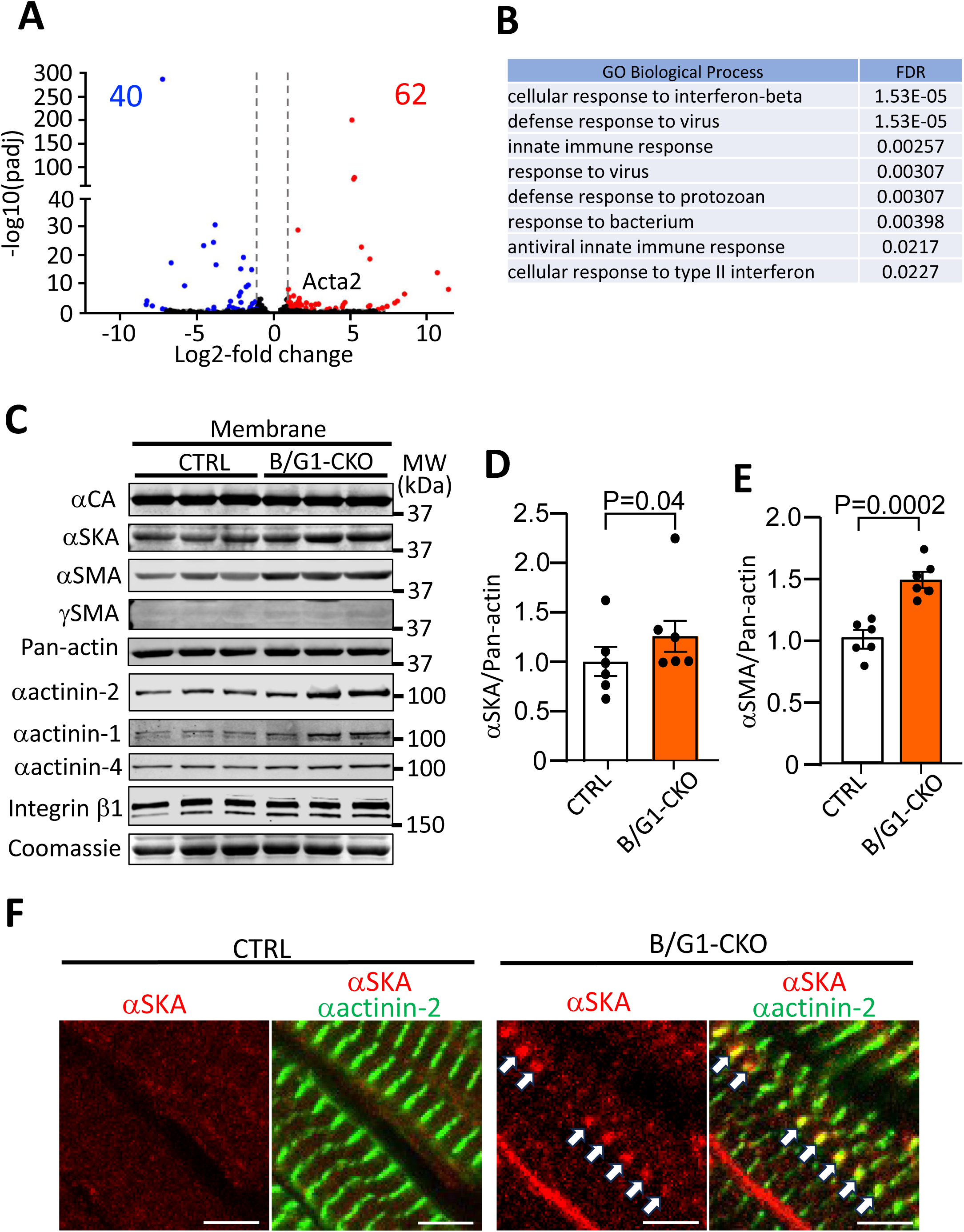
Characterization of heart-specific *Actb* and *Actg1* gene deletion. **(A)** Volcano plot of bulk RNA sequencing data from B/G1-CKO mouse hearts at 2 weeks of age; blue and red dots indicate down- and up-regulated genes, respectively. **(B)** Gene Ontology (GO) enrichment analysis and categories of the 62 upregulated genes, showing false discovery rate (FDR). **(C)** Western blotting of cardiac α-actin (αCA), skeletal muscle α-actin (αSKA), smooth muscle α-actin (αSMA), smooth muscle γ-actin (γSMA), pan-actin, α-actinin isoforms, and integrin β1 in membrane protein fractions from hearts of B/G1-CKO mice at 8-10 weeks of age; Coomassie staining is shown as a loading control, and numbers indicate molecular weight (kDa). **(D** and **E)** Quantification of αSKA protein (D) and αSMA protein (E) from the western blots (n = 6 per group). Data are shown as scatter plots with bars indicating mean ± SEM. **(F)** Representative immunohistochemical images of αSKA (red) and α-actinin-2 (green, cardiomyocyte marker) on heart sections from *Actb/g1^+/+-Myh6-Cre^*(control) or *Actb/g1^fl/fl-Myh6-Cre^* (B/G1-CKO) mice at 16 weeks of age. Scale bar, 5 µm. White arrows indicate accumulation of αSKA at the costamere.

Bulk RNA-seq also revealed significant upregulation of *Acta2*, encoding smooth muscle α-actin (αSMA), in B/G1-CKO hearts relative to controls (**Figure 2A** and **2C**, and **Supplementary Table 1**). Western blotting of membrane-enriched protein extracts confirmed increased expression of both skeletal α-actin (αSKA) and αSMA in B/G1-CKO hearts versus controls (**Figure 2C-E**). We next examined the subcellular localization of αSKA protein by immunohistochemistry (**Figure 2F**). αSKA was undetectable in the adult control hearts, whereas in B/G1-CKO hearts αSKA was expressed in a sarcomeric pattern that colocalized with α-actinin-2 at the Z-line, at the level of the costameres. Consistent with this compensatory remodeling, we also observed increased expression of both sarcomeric α-actinin-2 and non-sarcomeric α-actinin-1 in membrane protein extracts from B/G1-CKO hearts versus controls (**Figure 2C**). Because α-actinin proteins cross-link and stabilize actin filaments, these data suggest a compensatory attempt to reinforce the sarcolemma of B/G1-CKO cardiomyocytes.

### Cardiomyocyte-specific loss of cytoplasmic actin stabilizes the sarcolemma in dystrophin-deficient hearts

We next directly evaluated cardiac sarcolemmal stability in B/G1-CKO hearts by crossing this line into the *mdx* mouse model (**Figure 3A**). The *mdx* mouse carries a mutation in the Duchenne muscular dystrophy (*Dmd*) gene that when mutant results in sarcolemmal membrane instability in striated muscle [3,4]. This group and 3 other control genotype groups are shown in **Figures 3B** and **3C**. To assess membrane integrity across these groups, we performed Evans blue dye (EBD) uptake assays at 8 weeks of age (**Figure 3C-3E, Supplementary Figure 1**). As expected, *mdx* hearts showed increased EBD uptake relative to control hearts, but this membrane instability was lost when *Actb* and *Actg1* were also deleted with αMHC-Cre. These data show that B/G1-CKO hearts have greater membrane stability on the *mdx* background, supporting a primary role for the cytoplasmic actin network in regulating cardiomyocyte membrane properties.

**Figure 3.**
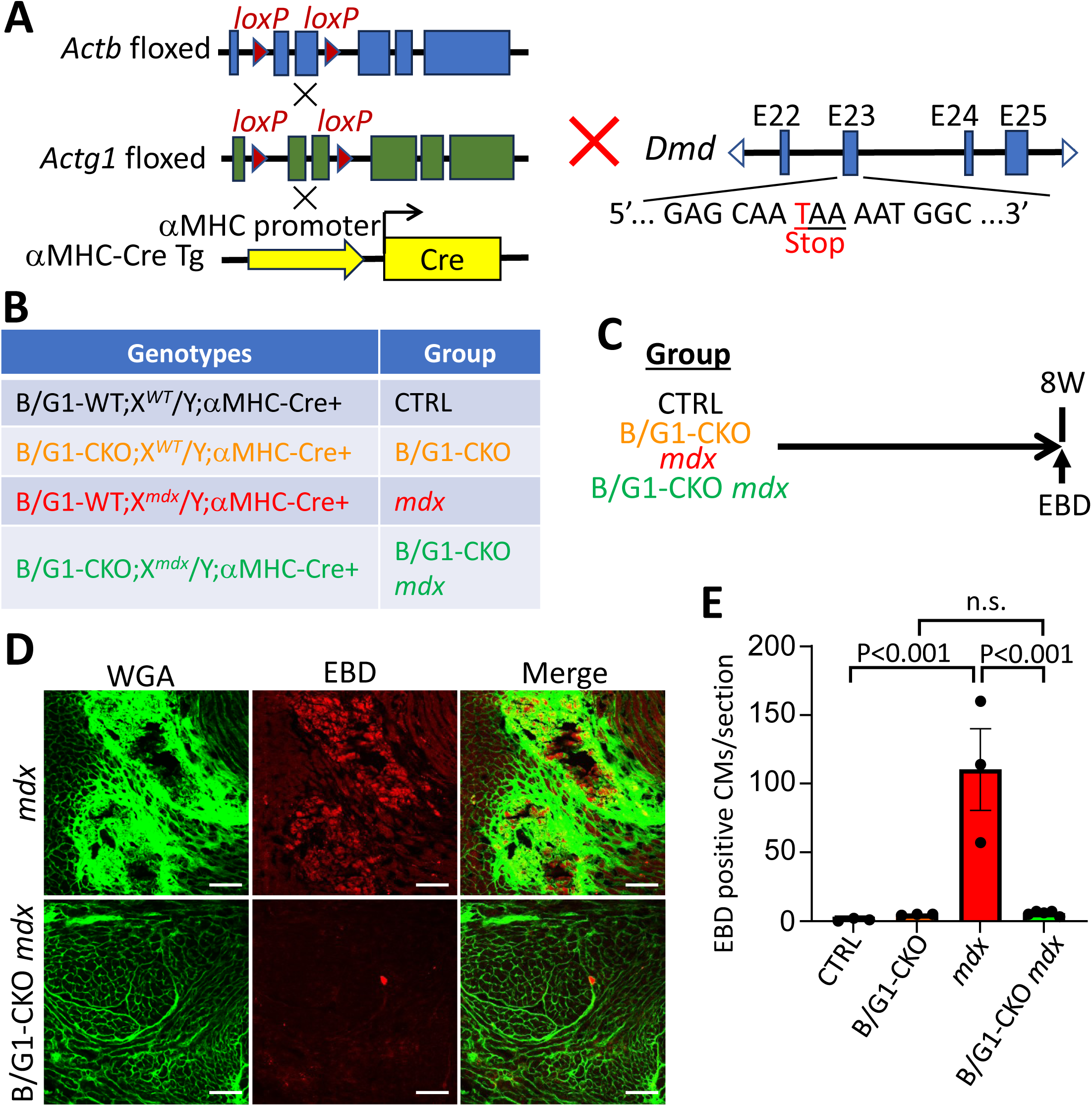
Heart-specific *Actb*/*Actg1*-deleted mice crossed with *mdx* mice. **(A)** B/G1-CKO mice were crossed with *mdx* mice to assess membrane stability. **(B)** Genotypes and groups generated from this cross: *Actb/g1^+/+-Myh6-Cre^*×*wt* (CTRL), *Actb/g1^fl/fl-Myh6-Cre^*×*wt* (B/G1-CKO), *Actb/g1^+/+-Myh6-Cre^*×*mdx* (MDX), and *Actb/g1^fl/fl-Myh6-Cre^*×*mdx* (B/G1-CKO *mdx*). **(C)** Experimental design used to assess membrane stability by Evans blue dye (EBD) uptake at 8 weeks of age (8W) in the 4 groups of mice. **(D)** Representative immunohistochemical images of wheat germ agglutinin (WGA) for membrane staining (green) and EBD for membrane integrity (red) in *mdx* (top) and B/G1-CKO *mdx* (bottom) heart sections. Scale bar, 100 µm. **(E)** Quantification of EBD uptake (n = 3 in control, B/G1-CKO, and *mdx* groups; n = 5 in B/G1-CKO MDX group). Data are shown as scatter plots with bars indicating mean ± SEM.

### Cytoplasmic actin deletion protects the heart from pressure overload-induced hypertrophy and dysfunction

The progressive pathogenesis of the failing heart is also linked to a weakened sarcolemma [30]. We therefore subjected control and B/G1-CKO adult mice to chronic pressure overload by transverse aortic constriction (TAC) over an 8-week time course (**Figure 4A**). TAC reduced ejection fraction (EF%) in control mice over 8 weeks, and this decline was partially prevented in B/G1-CKO hearts (**Figure 4B**). The marked cardiac hypertrophy observed in control mice after TAC was similarly attenuated in B/G1-CKO mice over the same time course (**Figure 4C**), and survival during the TAC procedure was greater in B/G1-CKO mice than in controls (**Figure 4D**).

**Figure 4.**
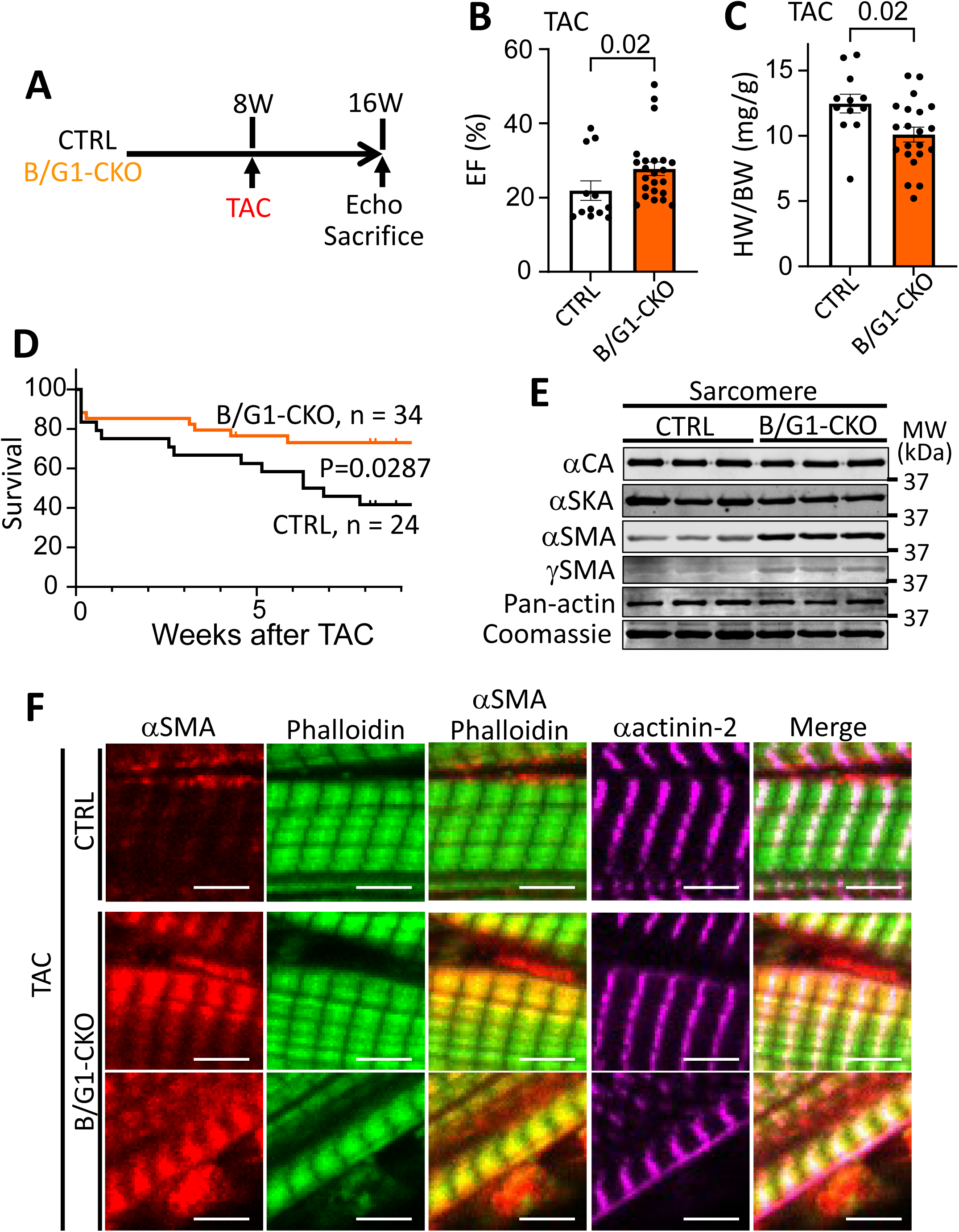
Pressure overload in heart-specific *Actb*/*Actg1*-deleted mice by transverse aortic constriction (TAC). **(A)** Experimental design of mice with TAC surgery at 8 weeks of age and then assessed for at 16 weeks of age (8 weeks post-surgery) by echocardiography, HW/BW ratio, Western blotting, and immunohistochemistry. **(B** and **C)** EF% (B) and HW/BW ratio (C) at 8 weeks after surgery (n = 12 control, n = 21 B/G1-CKO). Data are shown as scatter plots with bars indicating mean ± SEM. **(D)** Survival analysis after TAC surgery in the 2 groups of mice over the 8 weeks of TAC. **(E)** Western blots of cardiac α-actin (αCA), skeletal muscle α-actin (αSKA), smooth muscle α-actin (αSMA), smooth muscle γ-actin (γSMA), and pan-actin in the sarcomere protein fraction of B/G1-CKO hearts at 8 weeks after TAC; Coomassie staining is shown as a loading control, and numbers indicate molecular weight (kDa). **(F)** Representative immunohistochemistry from the heart histological sections of the indicated 2 groups of mice for αSMA (red), phalloidin for sarcomeric actin thin filaments (green), and α-actinin-2 (magenta). Scale bar, 4 µm.

In skeletal muscle, the cytoplasmic filamentous actin network is closely associated with costameres and links to the sarcomere [15,16]. We therefore examined actin isoform levels in the sarcomeric protein fraction of B/G1-CKO and control hearts at baseline (**Figure 4E**). αSKA protein levels in the sarcomere were unchanged in B/G1-CKO hearts relative to controls, but both αSMA and γSMA (encoded by *Actg2*) were induced and detectable at low levels in the sarcomere (**Figure 4E**). We extended this analysis to TAC-stimulated hearts using immunohistochemistry, which showed that αSMA protein was incorporated into the sarcomeric actin thin filament network, coincident with phalloidin staining, as well as localized to the costameric membrane attachment region (**Figure 4F**). These data indicate that re-expression of αSMA protein during disease stimulation can result in partial sarcomeric incorporation, an effect observed only with TAC stimulation and only in the B/G1-CKO background.

### Selective incorporation of induced actin isoforms into the sarcomere

Finally, of the 375 amino acids comprising each of the six mammalian actin isoforms, only 3-4 residues differ [10], although it remains uncertain if these modest changes underlie different functions across the 6 gene isoforms, or if regulation is based on differential expression of each of these genes. Given the induction of αSMA and αSKA at both the sarcomere and costamere with disease stimulation, we asked whether cytoplasmic β- and γ-actin were similarly induced within the sarcomere. To more directly test this we generated transgenic mice expressing both *Actb* and *Actg1* under control of the αMHC promoter using an inducible, tetracycline-repressor-based bicistronic dual-transgenic approach (icB/G1 Tg mice, **Supplementary Figure 2A**). We generated two founder lines that, in the absence of tetracycline or doxycycline, expressed the transgene in the heart, although only β-actin protein was significantly increased, suggesting that the IRES-based approach did not successfully drive γ-actin expression in this construct (**Supplementary Figure 2B-2D**). Overexpression of β-actin protein in the heart of these transgenic mice had no pathologic effect on heart weight normalized to body weight or on echocardiographic measured ejection fraction in adulthood (**Supplementary Figure 2E and 2F**). We also subjected icB/G1 Tg mice to TAC stimulation at 8 weeks of age for an additional 8 weeks but observed no change in the degree of cardiac hypertrophy or loss of ejection fraction relative to controls (**Supplementary Figure 2G-2I**). However, constitutive overexpression of *Actb* in the developing and adult heart resulted in β-actin protein being prominently incorporated into the cardiac sarcomeric thin filaments on either side of the α-actinin-labeled Z-line, but not within the subsarcolemmal cytoplasmic compartment (**Supplementary Figure 2J and 2K**). While this strategy, intended to ectopically re-induce the cytoplasmic actin filamentous network in the heart, did not achieve that specific goal, the results are nonetheless informative and show that simply increasing the β-actin content within the cardiac sarcomere was not itself pathological or beneficial.

## Discussion

This study identifies a previously unappreciated adaptive response of the cardiomyocyte to loss of its cytoplasmic actin network: rather than producing the myopathic, membrane-destabilizing phenotype reported after equivalent *Actb*/*Actg1* deletion in skeletal muscle [15,16], cardiomyocyte-specific deletion of cytoplasmic β- and γ-actin was well tolerated at baseline. Loss of this cytoplasmic actin network also triggered compensatory induction of the sarcomeric skeletal (αSKA) and smooth muscle (αSMA) α-actin isoforms at the sarcolemma and costamere. Overall B/G1-CKO hearts resisted membrane instability when combined with the dystrophin-deficient *mdx* mutation, and were substantially protected from hypertrophy, systolic dysfunction, and mortality after pressure overload. However, our results do not distinguish the mechanism whereby loss of cytoplasmic β- and γ-actin is protective to the sarcolemma in the heart with disease stimulation, as re-expression of these 2 actin isoforms during disease might help facilitate membrane remodeling that ultimately weakens the membrane, or the compensatory induction of the αSKA and αSMA isoforms at the level of costameres might be stabilizing to the membrane.

The tissue-specific divergence between our findings in heart and the published skeletal muscle data are notable. Muscle-specific deletion of *Actg1* or *Actb* produces progressive myofiber necrosis and costamere disorganization in skeletal muscle [15,16], and combining *Actg1* deletion with the *mdx* mutation neither worsened nor rescued the dystrophic phenotype, suggesting that γ-actin and dystrophin act within a shared pathway in that tissue [17]. In striking contrast, we found that cardiomyocyte-specific *Actb*/*Actg1* deletion actively protected the *mdx* heart from membrane instability. One explanation for this divergence is that cardiomyocytes, unlike skeletal myofibers, mount a robust compensatory transcriptional response to cytoplasmic actin loss, evident in our RNA-sequencing data as induction of *Acta2* and an interferon/innate-immune gene signature, the latter of which can protect the membrane from pathogen entry and otherwise provide a stabilizing influence. Alternatively, it is possible that induction of cytoplasmic β- and γ-actin with acute or chronic disease stimulation facilitates remodeling and instability of the focal adhesion and integrin complexes, as they could be processed for greater recycling as suggested previously [31].

Our attempt to directly re-establish the cytoplasmic actin network in the heart with the *Actb*/*Actg1* transgene was only partially successful: β-actin, but not γ-actin, was reliably overexpressed, and the overexpressed β-actin protein was incorporated into the sarcomere rather than the subsarcolemmal compartment. While this outcome limited our ability to test our initial hypothesis, it independently demonstrates that increasing β-actin content in the sarcomere did not produce a pathological phenotype at baseline or after pressure overload. This observation reinforces a broader observation that the cardiomyocyte tolerates considerable compositional flexibility in its actin network, whether through loss of the cytoplasmic network or through forced sarcomeric incorporation of a normally cytoplasmic actin isoform, without an obvious cost to baseline cardiac structure or function.

Costameres serve as the principal structural link between the sarcomere, the sarcolemma, and the extracellular matrix, and are increasingly recognized as sites of mechanotransduction that couple mechanical load to intracellular signaling in the cardiomyocyte [30]. The upregulation of α-actinin-1 and α-actinin-2, together with αSKA, at the membrane fraction of B/G1-CKO hearts is consistent with a coordinated strengthening of this costameric complex, analogous to the cardioprotective effect previously reported for costameric integrin overexpression during ischemia-reperfusion injury [7]. Considered together with the well-established role of sarcolemmal weakening in dystrophic and failing hearts [3–5,7–9], our data support a model in which the cardiomyocyte cytoskeleton is not a static structure but rather a dynamically regulated system that can be reinforced, whether by native compensatory remodeling or by therapeutic intervention, to resist membrane injury and hemodynamic stress.

### Conclusions and limitations

As a limitation, while our data show that αSKA and αSMA induction correlates with sarcolemmal protection, our approach cannot fully separate the specific contribution of this effect from the possibility that re-expression of cytoplasmic β- and γ-actin in the heart with disease stimulation is maladaptive, possibly by reducing focal adhesion complex stability. To address this limitation future genetic approaches that selectively prevent induction of *Acta1* or *Acta2* in the B/G1-CKO background will be needed to establish causality more directly. As another limitation, our inducible *Actb*/*Actg1* transgenic approach did not achieve subsarcolemmal re-expression of the cytoplasmic actin network, leaving open the question of whether restoring this network, could be detrimental to the heart, such as by destabilizing the focal adhesion complex. Despite these limitations, our findings establish that cardiomyocytes re-expression of the cytoplasmic β- and γ-actin isoforms in the heart is maladaptive and reduces sarcolemmal stability, suggesting that strategies to inhibit expression or activity of these 2 actin isoforms could be a therapeutic approach in forms of chronic heart disease resulting in failure.

## Supplementary Materials

Two Supplementary Tables and two Supplementary Figures

## Author Contributions

Conceptualization, Y.K., K.M.G. and J.D.M.; Methodology, Y.K., K.M.G., S-C.J.L., J.M.E., and J.D.M.; Validation, Y.K. and J.D.M.; Formal Analysis, Y.K.; Data Curation, Y.K., K.M.G., C.K., E.A., S. L.K.B., and S-C.J.L.; Writing – Original draft preparation, Y.K. and J.D.M.; Writing – Review and Editing, Y.K. and J.D.M.; Supervision, Y.K. and J.D.M.; Funding Acquisition: Y.K. and J.D.M.

## Institutional Review Board Statement

Not applicable (no human subjects or materials used).

## Informed Consent Statement

Not applicable.

## Data Availability Statement

The raw RNA-seq expression data were submitted to the GEO with an accession number of GSE339442 (embargoed until publication acceptance). All other data are contained within the manuscript or the supplements.

## Acknowledgements

We appreciate support from the Division of Comparative Medicine at CCHMC and equipment made available through the Cincinnati Children’s Research Foundation.

## References

1. Martin, S.S.; Aday, A.W.; Almarzooq, Z.I.; Anderson, C.A.M.; Arora, P.; Avery, C.L.; Baker-Smith, C.M.; Barone Gibbs, B.; Beaton, A.Z.; Boehme, A.K.;, et al. 2024 Heart disease and stroke statistics: a report of US and global data from the American Heart Association. Circulation 2024, 149, e347–e913, doi:10.1161/CIR.0000000000001209.

2. Dirkx, E.; da Costa Martins, P.A.; De Windt, L.J. Regulation of fetal gene expression in heart failure. Biochimica et Biophysica Acta (BBA) - Molecular Basis of Disease 2013, 1832, 2414–2424, doi:10.1016/j.bbadis.2013.07.023.

3. Yasuda S, Townsend D, Michele DE, Favre EG, Day SM, Metzger JM. Dystrophic heart failure blocked by membrane sealant poloxamer. Nature. 2005 Aug 18;436(7053):1025–9. doi: 10.1038/nature03844.

4. Johnstone, V.P.A.; Viola, H.M.; Hool, L.C. Dystrophic Cardiomyopathy—Potential role of calcium in pathogenesis, treatment and novel therapies. Genes 2017, 8, 108, doi:10.3390/genes8040108.

5. Turner, P.R.; Schultz, R.; Ganguly, B.; Steinhardt, R.A. Proteolysis results in altered leak channel kinetics and elevated free calcium in mdx muscle. J Membr Biol 1993, 133, 243–251, doi:10.1007/BF00232023.

6. Burr AR, Molkentin JD. Genetic evidence in the mouse solidifies the calcium hypothesis of myofiber death in muscular dystrophy. Cell Death Differ. 2015 Sep;22(9):1402–12. doi: 10.1038/cdd.2015.65.

7. Okada, H.; Lai, N.C.; Kawaraguchi, Y.; Liao, P.; Copps, J.; Sugano, Y.; Okada-Maeda, S.; Banerjee, I.; Schilling, J.M.; Gingras, A.R.;, et al. Integrins protect cardiomyocytes from ischemia/reperfusion injury. J Clin Invest 2013, 123, 4294–4308, doi:10.1172/JCI64216.

8. Lynch JM, Maillet M, Vanhoutte D, Schloemer A, Sargent MA, Blair NS, Lynch KA, Okada T, Aronow BJ, Osinska H, Prywes R, Lorenz JN, Mori K, Lawler J, Robbins J, Molkentin JD. A thrombospondin-dependent pathway for a protective ER stress response. Cell. 2012 Jun 8;149(6):1257–68. doi: 10.1016/j.cell.2012.03.050.

9. Vanhoutte D, Schips TG, Kwong JQ, Davis J, Tjondrokoesoemo A, Brody MJ, Sargent MA, Kanisicak O, Yi H, Gao QQ, Rabinowitz JE, Volk T, McNally EM, Molkentin JD. Thrombospondin expression in myofibers stabilizes muscle membranes. Elife. 2016 Sep 26;5:e17589. doi: 10.7554/eLife.17589.

10. Perrin, B.J.; Ervasti, J.M. The actin gene family: function follows isoform. Cytoskeleton 2010, 67, 630–634, doi:10.1002/cm.20475.

11. Heissler SM, Chinthalapudi K. Structural and functional mechanisms of actin isoforms. FEBS J. 2025 Feb;292(3):468–482. doi: 10.1111/febs.17153.

12. Kumar, A.; Crawford, K.; Close, L.; Madison, M.; Lorenz, J.; Doetschman, T.; Pawlowski, S.; Duffy, J.; Neumann, J.; Robbins, J.;, et al. Rescue of cardiac alpha-actin-deficient mice by enteric smooth muscle gamma-actin. Proc Natl Acad Sci USA 1997, 94, 4406–4411, doi:10.1073/pnas.94.9.4406.

13. Grimes KM, Maillet M, Swoboda CO, Bowers SLK, Millay DP, Molkentin JD. MEK1-ERK1/2 signaling regulates the cardiomyocyte non-sarcomeric actin cytoskeletal network. Am J Physiol Heart Circ Physiol. 2024 Jan 1;326(1):H180–H189. doi: 10.1152/ajpheart.00612.2023.

14. Mehdiabadi NR, Boon Sim C, Phipson B, Kalathur RKR, Sun Y, Vivien CJ, Ter Huurne M, Piers AT, Hudson JE, Oshlack A, Weintraub RG, Konstantinov IE, Palpant NJ, Elliott DA, Porrello ER. Defining the fetal gene program at single-cell resolution in pediatric dilated cardiomyopathy. Circulation. 2022 Oct 4;146(14):1105–1108. doi: 10.1161/CIRCULATIONAHA.121.057763..

15. Sonnemann, K.J.; Fitzsimons, D.P.; Patel, J.R.; Liu, Y.; Schneider, M.F.; Moss, R.L.; Ervasti, J.M. Cytoplasmic gamma-actin is not required for skeletal muscle development but its absence leads to a progressive myopathy. Dev Cell 2006, 11, 387–397, doi:10.1016/j.devcel.2006.07.001.

16. Prins, K.W.; Call, J.A.; Lowe, D.A.; Ervasti, J.M. Quadriceps myopathy caused by skeletal muscle-specific ablation of β(cyto)-actin. J Cell Sci 2011, 124, 951–957, doi:10.1242/jcs.079848.

17. Prins, K.W.; Lowe, D.A.; Ervasti, J.M. Skeletal muscle-specific ablation of γ(cyto)-actin does not exacerbate the mdx phenotype. PLoS ONE 2008, 3, e2419, doi:10.1371/journal.pone.0002419.

18. Perrin, B.J.; Sonnemann, K.J.; Ervasti, J.M. β-actin and γ-actin are each dispensable for auditory hair cell development but required for stereocilia maintenance. PLOS Genetics 2010, 6, e1001158, doi:10.1371/journal.pgen.1001158.

19. Belyantseva, I.A.; Perrin, B.J.; Sonnemann, K.J.; Zhu, M.; Stepanyan, R.; McGee, J.; Frolenkov, G.I.; Walsh, E.J.; Friderici, K.H.; Friedman, T.B.;, et al. γ-actin is required for cytoskeletal maintenance but not development. Proceedings of the National Academy of Sciences 2009, 106, 9703–9708, doi:10.1073/pnas.0900221106.

20. Kuwabara, Y.; York, A.J.; Lin, S.-C.; Sargent, M.A.; Grimes, K.M.; Pirruccello, J.P.; Molkentin, J.D. A human FLII gene variant alters sarcomeric actin thin filament length and predisposes to cardiomyopathy. Proceedings of the National Academy of Sciences 2023, 120, e2213696120, doi:10.1073/pnas.2213696120.

21. Minerath, R.A.; Kasam, R.K.; Swoboda, C.O.; Prasad, V.; Grimes, K.M.; Blair, N.S.; Khalil, H.; Alfieri, C.M.; Eads, L.; Saviola, A.J.;, et al. Cardiomyocyte-expressed TGFβ signals to fibroblasts to program early heart maturation and adult myocyte identity. 2025, BioRxiv 2025.09.22.677845. doi: 10.1101/2025.09.22.677845.

22. Sanbe A, Gulick J, Hanks MC, Liang Q, Osinska H, Robbins J. Reengineering inducible cardiac-specific transgenesis with an attenuated myosin heavy chain promoter. Circ Res. 2003 Apr 4;92(6):609–16. doi: 10.1161/01.RES.0000065442.64694.9F.

23. Brito-Estrada, O.; Kuwabara, Y.; Gibson, A.M.; Hassel, K.R.; Kamradt, M.L.; Verry, J.P.; Blair, N.S.; Bround, M.J.; Huo, J.; Molkentin, J.D.;, et al. DWORF gene therapy improves cardiac calcium handling and mitochondrial function. Circulation Research 2025, 137, 1072–1088, doi:10.1161/CIRCRESAHA.125.326550.

24. Sert, N.P. du; Hurst, V.; Ahluwalia, A.; Alam, S.; Avey, M.T.; Baker, M.; Browne, W.J.; Clark, A.; Cuthill, I.C.; Dirnagl, U.;, et al. The ARRIVE guidelines 2.0: updated guidelines for reporting animal research. PLOS Biology 2020, 18, e3000410, doi:10.1371/journal.pbio.3000410.

25. Kuwabara, Y.; Keezer, C.; Lin, S.-C.J.; Rajput, A.; Molkentin, J.D. Heart-Specific and Conditional Deletion of the Immt Gene Reveals Its Role in Regulating Mitochondrial Structure and Total Heart Metabolism. Cells 2026, 15, 505, doi:10.3390/cells15060505.

26. Vanhoutte, D.; Schips, T.G.; Minerath, R.A.; Huo, J.; Kavuri, N.S.S.; Prasad, V.; Lin, S.-C.; Bround, M.J.; Sargent, M.A.; Adams, C.M.;, et al. Thbs1 regulates skeletal muscle mass in a TGFβ-Smad2/3-ATF4-dependent manner. Cell Reports 2024, 43, 114149, doi:10.1016/j.celrep.2024.114149.

27. Grimes, K.M.; Barefield, D.Y.; Kumar, M.; McNamara, J.W.; Weintraub, S.T.; de Tombe, P.P.; Sadayappan, S.; Buffenstein, R. The naked mole-rat exhibits an unusual cardiac myofilament protein profile providing new insights into heart function of this naturally subterranean rodent. Pflugers Arch 2017, 469, 1603–1613, doi:10.1007/s00424-017-2046-3.

28. Kuwabara, Y.; Tsuji, S.; Nishiga, M.; Izuhara, M.; Ito, S.; Nagao, K.; Horie, T.; Watanabe, S.; Koyama, S.; Kiryu, H.;, et al. Lionheart LincRNA alleviates cardiac systolic dysfunction under pressure overload. Commun Biol 2020, 3, 434, doi:10.1038/s42003-020-01164-0.

29. Davis, J.; Maillet, M.; Miano, J.M.; Molkentin, J.D. Lost in transgenesis: a user’s guide for genetically manipulating the mouse in cardiac research. Circ Res 2012, 111, 761–777, doi:10.1161/CIRCRESAHA.111.262717.

30. Samarel, A.M. Costameres, focal adhesions, and cardiomyocyte mechanotransduction. Am J Physiol Heart Circ Physiol 2005, 289, H2291–H2301, doi:10.1152/ajpheart.00749.2005.

31. Vicente-Manzanares M, Choi CK, Horwitz AR. Integrins in cell migration--the actin connection. J Cell Sci. 2009 Jan 15;122(Pt 2):199–206. doi: 10.1242/jcs.018564.

