## Supplementary Figures and Tables for "Deletion of cytoplasmic β/γ-actin in the mouse heart protects from disease by augmenting sarcolemma stability"

### Supplementary Fig. 1

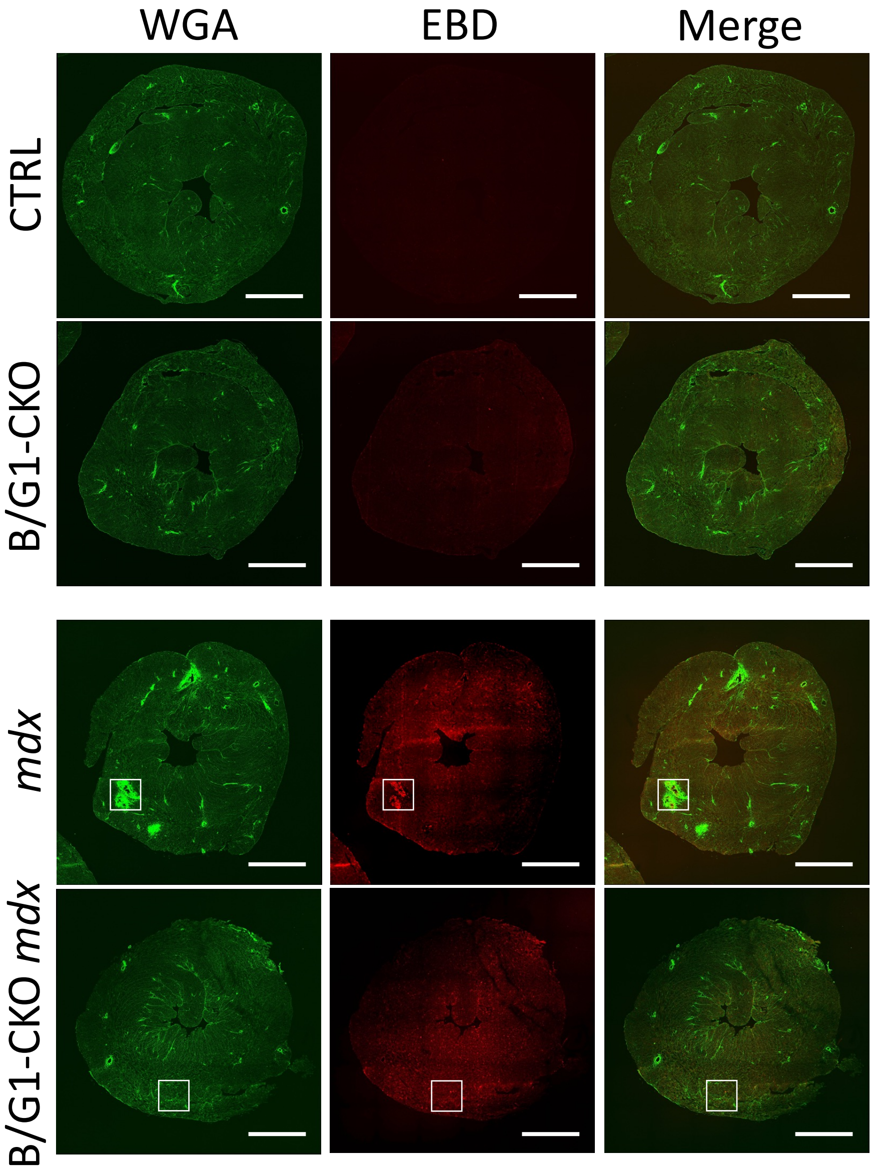


**Supplementary Figure 1. Membrane stability assessment by EBD uptake in B/G1-CKO mice crossed with *mdx* mice.** Representative whole heart cross-section histological images of wheat germ agglutinin (WGA) for membrane staining (green) and Evans blue dye (EBD) for membrane integrity (red) on Control (top row), B/G1-CKO (2nd row), *mdx* (3rd row), and B/G1-CKO *mdx* (bottom row) at 8 weeks of age. Scale bar, 1 mm. White square indicates the area shown in Figure 3D.

### Supplementary Fig. 2

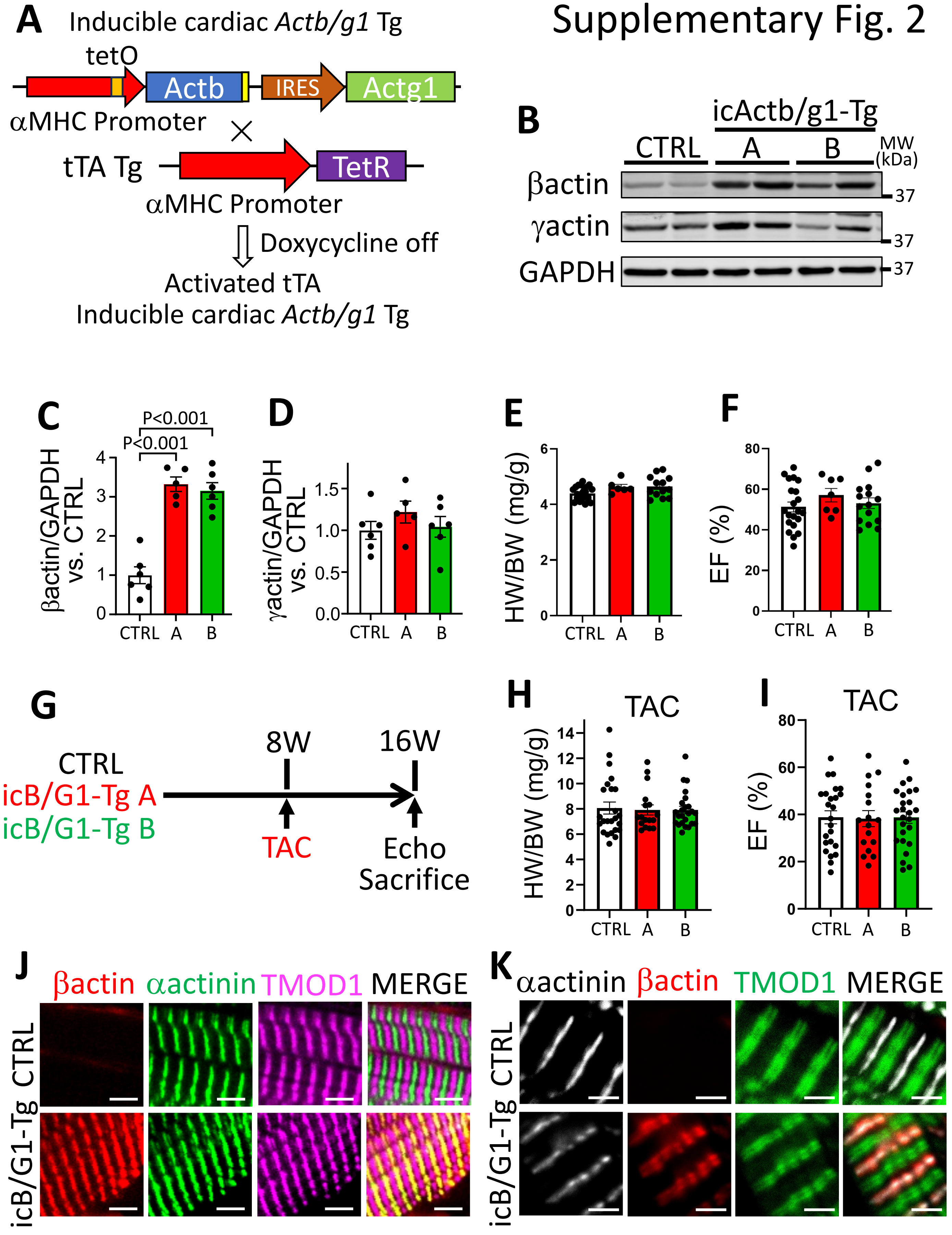

**Supplementary Figure 2. Cardiomyocyte-specific, inducible *Actb*/*Actg1* transgenic mice. (A)** Strategy for generating the inducible cardiac *Actb*/*Actg1* transgenic mouse (icActb/g1-Tg or icB/G1-Tg) using a tetracycline-off system driven by the bigenic αMHC promoter promoter system that is shut down in the presence of doxycycline, or activated constitutively if double transgenic mice are never given doxycycline. **(B)** Western blots of β-actin and γ-actin in whole-heart protein extracts from icActb/g1-Tg mice; GAPDH served as a loading control, and numbers indicate molecular weight (kDa) in control mice and 2 different transgenic lines as A or B. **(C** and **D)** Quantification of β-actin (C) and γ-actin (D) by western blotting in icActb/g1-Tg mouse hearts versus control (n = 6 per group). **(E)** HW/BW ratio at 16 weeks of age (n = 21 control, n = 6 icActb/g1-Tg A, n = 13 icActb/g1-Tg B). **(F)** EF% by echocardiography at 16 weeks of age (n = 21 control, n = 7 icActb/g1-Tg A, n = 15 icActb/g1-Tg B). **(G)** Experimental time-line for pressure overload by TAC in 8 week old in icB/G1-Tg or control mice, followed by phenotyping at 16 weeks of age (8 weeks post-surgery). **(H)** HW/BW ratio at 8 weeks after TAC (n = 24 control, n = 17 icB/G1-Tg A, n = 24 icB/G1-Tg B). **(I)** EF% at 8 weeks after TAC (n = 24 control, n = 17 icB/G1-Tg A, n = 25 icB/G1-Tg B). Data in C-F, H, and I are shown as scatter plots with bars indicating mean ± SEM. **(J)** Representative immunohistochemical confocal images from heart histological sections from the 2 groups of mice shown for β-actin (red), α-actinin-2 (green, Z-disc marker), and TMOD1 (magenta, marker of the pointed actin thin filament end near the M-line) at 16 weeks of age. tTA transgene only represented the control line. Scale bar, 4 µm. **(K)** Representative immunohistochemical images of α-actinin-2 (white), β-actin (red), and TMOD1 (green) on histological heart sections from tTA (control) or icB/G1-Tg mice at 16 weeks of age, imaged by super-resolution confocal microscopy. Scale bar, 2 µm.

### Supplementary Table 1. Upregulated Genes in B/G1-CKO mouse heart

| | Gene_name | log2FoldChange | padj | Gene_description |
|---|---|---|---|---|
| 1 | Gbp2b | 11.39520038 | 2.49609E-09 | guanylate binding protein 2b |
| 2 | Gm20400 | 10.67732258 | 3.75063E-15 | predicted gene 20400 |
| 3 | Gm12840 | 8.556162222 | 1.12564E-07 | predicted gene 12840 |
| 4 | Zfp459 | 8.086743612 | 2.93723E-05 | zinc finger protein 459 |
| 5 | BC018473 | 7.923913033 | 0.000173374 | cDNA sequence BC018473 |
| 6 | Trac | 7.423996306 | 0.002439024 | T cell receptor alpha constant |
| 7 | Gm32171 | 7.420566835 | 0.002826799 | predicted gene, 32171 |
| 8 | Fam220-ps | 7.107833821 | 0.00859294 | family with sequence similarity 220, pseudogene |
| 9 | Hist1h2al | 6.838856951 | 0.03003564 | histone cluster 1, H2al |
| 10 | Gm16092 | 6.313261512 | 7.07398E-20 | predicted gene 16092 |
| 11 | Gm18562 | 6.307205903 | 0.001135777 | predicted gene, 18562 |
| 12 | Tgtp1 | 6.229108551 | 0.002636734 | T cell specific GTPase 1 |
| 13 | Gm19277 | 5.763858247 | 4.60653E-24 | predicted gene, 19277 |
| 14 | Rps3a3 | 5.309074059 | 5.12043E-68 | ribosomal protein S3A3 |
| 15 | Rps3a2 | 5.262301866 | 1.05205E-64 | ribosomal protein S3A2 |
| 16 | Gm15459 | 5.146647603 | 1.8133E-188 | predicted gene 15459 |
| 17 | Gm26825 | 4.696884499 | 0.000103572 | predicted gene, 26825 |
| 18 | Zbp1 | 4.573963336 | 0.000669904 | Z-DNA binding protein 1 |
| 19 | Ifi208 | 4.530167327 | 0.005337162 | interferon activated gene 208 |
| 20 | Cd300lf | 4.106116761 | 0.010306046 | CD300 molecule like family member F |
| 21 | Trim34b | 3.699488115 | 0.018260735 | tripartite motif-containing 34B |
| 22 | H2-Q5 | 3.538501139 | 0.02704737 | histocompatibility 2, Q region locus 5 |
| 23 | Gm5900 | 2.989823065 | 0.000381824 | predicted pseudogene 5900 |
| 24 | Gm8399 | 2.795567517 | 0.000486326 | predicted gene 8399 |
| 25 | Gm12250 | 2.68109226 | 0.03003564 | predicted gene 12250 |
| 26 | Zfp599 | 2.538769184 | 0.043701489 | zinc finger protein 599 |
| 27 | Gbp6 | 2.497561195 | 0.000929461 | guanylate binding protein 6 |
| 28 | Ifit3 | 2.199204487 | 0.000150403 | interferon-induced protein with tetratricopeptide repeats 3 |
| 29 | Gm8325 | 2.170527937 | 0.00873987 | predicted pseudogene 8325 |
| 30 | Pdk4 | 1.887385731 | 0.000829983 | pyruvate dehydrogenase kinase, isoenzyme 4 |
| 31 | Cracr2a | 1.8052636 | 0.000381824 | calcium release activated channel regulator 2A |
| 32 | Gm12715 | 1.753780846 | 0.000657382 | predicted gene 12715 |
| 33 | Angptl4 | 1.749149264 | 3.90881E-06 | angiopoietin-like 4 |
| 34 | H2-Ab1 | 1.736585273 | 0.028367992 | histocompatibility 2, class II antigen A, beta 1 |
| 35 | Atp6v0e2 | 1.724601441 | 6.24493E-05 | ATPase, H+ transporting, lysosomal V0 subunit E2 |
| 36 | Bdnf | 1.718388534 | 0.032709596 | brain derived neurotrophic factor |
| 37 | Gck | 1.67374167 | 4.8212E-05 | glucokinase |
| 38 | Actg1 | 1.650100578 | 4.6494E-30 | actin, gamma, cytoplasmic 1 |
| 39 | Asns | 1.6362544 | 0.002903071 | asparagine synthetase |
| 40 | Tctex1d2 | 1.548790879 | 0.005280998 | Tctex1 domain containing 2 |
| 41 | H2-Q4 | 1.492091516 | 0.000486326 | histocompatibility 2, Q region locus 4 |
| 42 | Gm6548 | 1.470150244 | 0.049134617 | predicted gene 6548 |
| 43 | Tap1 | 1.434187928 | 0.025382972 | transporter 1, ATP-binding cassette, sub-family B (MDR/TAP) |
| 44 | Sorl1 | 1.370397489 | 5.3647E-07 | sortilin-related receptor, LDLR class A repeats-containing |
| 45 | Klhdc7a | 1.306150704 | 0.017710468 | kelch domain containing 7A |
| 46 | Ifi27l2a | 1.301473196 | 0.03003564 | interferon, alpha-inducible protein 27 like 2A |
| 47 | Ttc38 | 1.266763693 | 0.001139042 | tetratricopeptide repeat domain 38 |
| 48 | Tmod4 | 1.244964725 | 0.009135204 | tropomodulin 4 |
| 49 | Gpr137b | 1.198033667 | 0.001139042 | G protein-coupled receptor 137B |
| 50 | Dtx3l | 1.194208042 | 0.001027871 | deltex 3-like, E3 ubiquitin ligase |
| 51 | Cybb | 1.186388909 | 0.038844142 | cytochrome b-245, beta polypeptide |
| 52 | Acot1 | 1.17332349 | 0.010981583 | acyl-CoA thioesterase 1 |
| 53 | Apobec3 | 1.171962399 | 0.00517384 | apolipoprotein B mRNA editing enzyme, catalytic polypeptide 3 |
| 54 | Mybpc2 | 1.169705679 | 0.00011413 | myosin binding protein C, fast-type |
| 55 | Gm3608 | 1.111172954 | 0.036194468 | predicted gene 3608 |
| 56 | Fbln1 | 1.084590989 | 0.001616809 | fibulin 1 |
| 57 | Acsf2 | 1.074682552 | 1.27923E-06 | acyl-CoA synthetase family member 2 |
| 58 | Hba-a1 | 1.058711376 | 3.56305E-05 | hemoglobin alpha, adult chain 1 |
| 59 | Ifit2 | 1.054036153 | 0.02259982 | interferon-induced protein with tetratricopeptide repeats 2 |
| 60 | Kbtbd12 | 1.053365248 | 6.96788E-05 | kelch repeat and BTB (POZ) domain containing 12 |
| 61 | Acta2 | 1.041231491 | 2.21768E-09 | actin, alpha 2, smooth muscle, aorta |
| 62 | Casq1 | 1.010970888 | 0.002059332 | calsequestrin 1 |

### Supplementary Table2. Downregulated Genes in B/G1-CKO mouse heart

| | Gene_name | log2FoldChange | padj | Gene_description |
|---|---|---|---|---|
| 1 | Klk1b22 | -8.1763071 | 0.00055057 | kallikrein 1-related peptidase b22 |
| 2 | AC131339.4 | -8.1190267 | 2.5706E-05 | novel transcript |
| 3 | Klra9 | -7.7726737 | 0.00161646 | killer cell lectin-like receptor subfamily A, member 9 |
| 4 | AC131339.2 | -7.1125167 | 2.971E-274 | novel transcript |
| 5 | Gm44366 | -7.1017099 | 0.01730348 | predicted gene, 44366 |
| 6 | Gm37612 | -6.9279301 | 0.03603248 | predicted gene, 37612 |
| 7 | Gm8741 | -6.5480651 | 1.5579E-18 | predicted gene 8741 |
| 8 | Slc15a2 | -5.689199 | 1.6268E-10 | solute carrier family 15 (H+/peptide transporter), member 2 |
| 9 | Kcnj14 | -4.4490811 | 1.3688E-24 | potassium inwardly-rectifying channel, subfamily J, member 14 |
| 10 | 5830444B04Rik | -4.1743198 | 0.04668478 | RIKEN cDNA 5830444B04 gene |
| 11 | AI506816 | -3.8191215 | 9.4355E-26 | expressed sequence AI506816 |
| 12 | Rps6ka6 | -3.7981758 | 0.04234119 | ribosomal protein S6 kinase polypeptide 6 |
| 13 | 2610035D17Rik | -3.7929675 | 0.005281 | RIKEN cDNA 2610035D17 gene |
| 14 | Gm6969 | -3.7091178 | 7.3754E-32 | predicted pseudogene 6969 |
| 15 | Rab6b | -3.6470718 | 7.306E-18 | RAB6B, member RAS oncogene family |
| 16 | Gm8188 | -2.9321164 | 0.04978708 | predicted gene 8188 |
| 17 | Opcml | -2.7883033 | 0.00243902 | opioid binding protein/cell adhesion molecule-like |
| 18 | Serpina3n | -2.7159061 | 3.9278E-05 | serine (or cysteine) peptidase inhibitor, clade A, member 3N |
| 19 | Cpa3 | -2.532185 | 0.00925238 | carboxypeptidase A3, mast cell |
| 22 | Arhgap36 | -2.3777888 | 0.01602319 | Rho GTPase activating protein 36 |
| 23 | Shisal1 | -2.2282681 | 0.0238435 | shisa like 1 |
| 24 | Art4 | -2.1573577 | 5.1644E-07 | ADP-ribosyltransferase 4 |
| 25 | Hcn1 | -2.0541613 | 2.4948E-08 | hyperpolarization-activated, cyclic nucleotide-gated K+ 1 |
| 26 | Ush1c | -2.0457795 | 1.8639E-16 | USH1 protein network component harmonin |
| 27 | Hdhd3 | -2.0389922 | 0.00010357 | haloacid dehalogenase-like hydrolase domain containing 3 |
| 28 | Elfn2 | -1.9174478 | 0.04913462 | leucine rich repeat and fibronectin type III, extracellular 2 |
| 29 | Rpl3-ps1 | -1.875621 | 1.7338E-20 | ribosomal protein L3, pseudogene 1 |
| 30 | Wdfy1 | -1.6998765 | 3.5777E-10 | WD repeat and FYVE domain containing 1 |
| 31 | Gramd1b | -1.5564143 | 7.3431E-11 | GRAM domain containing 1B |
| 32 | Mrap | -1.4840258 | 0.03003564 | melanocortin 2 receptor accessory protein |
| 33 | Gm40841 | -1.4567964 | 0.00635344 | predicted gene, 40841 |
| 34 | Gal3st2c | -1.4249293 | 0.0077107 | galactose-3-O-sulfotransferase 2C |
| 35 | Slc4a8 | -1.4097243 | 0.02705646 | solute carrier family 4 (anion exchanger), member 8 |
| 36 | Bmper | -1.4067597 | 0.00915483 | BMP-binding endothelial regulator |
| 37 | C1qtnf6 | -1.3451743 | 3.5074E-16 | C1q and tumor necrosis factor related protein 6 |
| 38 | Gsg1l | -1.2888853 | 0.00025375 | GSG1-like |
| 39 | Gm43305 | -1.2289146 | 0.03619447 | predicted gene 43305 |
| 40 | Slc2a12 | -1.1321276 | 0.02357521 | solute carrier family 2 (facilitated glucose transporter), member 12 |
| 41 | Alad | -1.1004156 | 5.0072E-05 | aminolevulinate, delta-, dehydratase |
| 42 | Lrrc4b | -1.0301642 | 0.01981648 | leucine rich repeat containing 4B |
